# Using eDNA methodologies to identify the presence/absence of *S. thermophilus* in yogurt products

**DOI:** 10.1101/414060

**Authors:** Vladimir Bejdo, Claire Gao

## Abstract

The advent of low-cost nuclear acid extraction allows for the creation of low-cost assays which can specifically be used to determine the presence or absence of bacteria in a variety of environments. Commercially sold dietary yogurt claims to contain bacteria forming a microbiome which has been previously linked to improved health outcomes in incidence rates of type 2 diabetes in vulnerable populations. The predicted outcome was that a main bacterial culture used in yogurt production *(S. thermophilus)* would be present and would be detectable using eDNA methodologies since it is fundamental to the making of yogurt. We gathered DNA from yogurt using yogurt dilutions and filter paper; we then extracted the DNA and also sequenced and used PCR to amplify the DNA. We sequenced PCR products to verify their identity through processing with publicly available BLAST tools which reference already accessioned bacterial genomes. Yogurt from four different commercially available brands (in the U.S.) was tested; not all yogurts tested positive for the bacteria, with higher concentrations of the bacteria in imported Greek-style yogurts, lower concentrations in domestically produced Greek-style yogurts, and no bacterial DNA detected in domestically produced ‘regular’ yogurt products. This research suggests that not all yogurts are equal, putting into question the whole-sale claims made on dietary yogurt’s probiotic preventative health effects and calling for a more detailed analysis to determine firm causal links between the microbiota of yogurts and preventative health effects.

## 1. Introduction

Previous studies have shown (and advertisers often promote) the positive health impacts which seem to be correlated to eating yogurt and other fermented foods. These effects have been attributed to the probiotics which promote the fermentative processes that create these products; in dietary yogurt, these two bacteria are *L. delbrueckii ssp. bulgaricus* and *S. thermophilus* [1].

Consuming yogurt on a regular basis has been shown to reduce the risk of T2D in elderly populations at a high risk of cardiovascular disease in Spain [2]. The same effect has been found in general adult populations in the United States, where over 194 thousand adults were examined on eating habits and in particular on the types of dairy that they ate in a study which conducted meta-analysis to find similar effects caused from yogurt consumption [3]. In order for these yogurt products to have any positive effect, they need to have the proper bacteria which facilitate fermentation; this study aims to introduce a methodology to prove that yogurt, a potential preventative against widespread disease like type 2 diabetes, as commercially sold, truly has the effects attributed to these bacteria that have been studied and publicized domestically and internationally by confirming their presence in the yogurt microbiome.

No recorded efforts have been made to confirm manufacturer claims on the presence of live cultures in yogurt products. Most studies in the past that have analyzed the bacterial content of yogurt have utilized phenotypic and fermentative profiling, which is highly variable for each strain of the bacteria potentially present [4, 5]. An emerging method of bacterial identification in environmental sources involves the use of DNA sequencing, or at the very least PCR/amplification. Some of the most prevalent methods for using DNA in bacterial identification involve DNA fingerprinting by testing through methods like random amplified polymorphic DNA (RAPD) analysis and restriction fragment-length polymorphism (RFLP) analysis [6, 7, 8, 9]–both methods that can produce results even when the specifics of the sample tested are not well known [10]. However, these methods may produce wildly variable PCR products, are at times resource-heavy, and are not always reproducible due to protocols that are extremely dependent on individual testing and laboratory conditions [11]. Due to these issues, these methods are sometimes not seen as stable enough to be used as stand-alone methods [12], and are often accompanied with existing profiling methods. DNA sequencing has, however, advanced to the point where we can create species specific primers for the particular and replicable am-plification of bacterial DNA samples, potentially negating the need for a hyper-general testing method. Methods recently used for the isolation of environmental DNA have become increasingly cost effective, costing a few cents per sample identified, and can provide quick and certain results [13].

To make use of this technology, we have conducted preliminary testing of commercially available dietary yogurts in order to begin to form a consistent, cost-effective methodology for the use of eDNA methods in the identification of probiotics in yogurt. This paper comes to preliminary conclusions and suggestions for further research to help protect trade and consumer interests by pointing out potential false advertising, and suggests that this methodology provides an opportunity to more definitively prove causation between these bacteria and the effects of yogurt on vulnerable groups by proving whether or not these bacteria are actually present in the disease-preventative yogurt noted in literature.

## 2. Methodology

This section outlines the methodology used in yogurt sample collection, DNA extraction, amplification, sequencing, and sequence processing.

In order to conduct this experiment, four different brands of commercially available yogurt products (available at time of publication in Seattle, WA) were used. The four yogurts tested were Fage’s *Total* (an imported Greek yogurt product), Chobani Plain Yogurt, Greek Gods Greek Yogurt (domestic Greek yogurt products), and Yoplait Original Strawberry (a domestic yogurt product). All the brands claim to have been produced using starter cultures containing both tested bacteria, and claim to have ‘live and active’ cultures.

### 2.1. Sample collection

Samples were prepared for collection by preparing a 1:25 dilution in a test tube; this was done by taking 0.5 mL of yogurt and adding it to 12 mL of distilled water. This mixture is then capped and agitated for about 30 seconds. Precipitation may be present; this does not affect the overall sample quality. For the actual collection process, a piece of standard Whatman Grade 1 (15mm diameter) qualitative filter paper is placed into the mixture for 30 seconds with forceps, retrieved, and stored in an appropriately sized (2 mL) screw capped microcentrifuge tube with 1 mL of NP-40 lysis buffer for storage. This process allows for the DNA to be isolated and quickly prepared for extraction shortly after sampling. Alternatively, samples from the solution can be pipetted onto the filter paper for storage; the ‘dipping’ method allows for less specialized tools to be used to achieve the same effect, and is the one used in this experiment.

### 2.2. DNA extraction/purification

DNA extraction/purification was performed with a method similar to the one found in the article *Nucleic acid purification from plants, animals and microbes in under 30 seconds* which calls for directly dipping a dipstick exposed to the sample to an amplification reaction after a few dips in a wash buffer [13]. Instead, to extract as much DNA as possible from samples and additionally keep material costs down, a small (roughly 1 cm x 1 cm large) piece of the original filter paper was placed into 500 μL Tris buffer and then incubated in 200 μL of distilled water in order to extract DNA off the filter paper. This allows for sufficient time to have a solution that can be used for amplification.

### 2.3. PCR amplification

The particular PCR beads used to perform PCR amplification were General Electric’s illustra™ PuReTaq™ Ready-To-Go™ PCR beads. The procedure is performed as standard–adding a reaction bead, 1 μL of each forward and reverse primer for the desired bacteria, 18 μL of water, and 5 μL of the DNA template prepared in the previous stage into a 200 μL Eppendorf tube.

### 2.4. Primers

Primers were designed using publicly available accessioned bacterial genomes for *S. thermophilus* using NCBI’s Primer-BLAST. The forward primer is GCT TTA GGG CTA GCG TCG AT, while the reverse primer is TAG GTC CCG ACT AAC CCA GG, listed in 5′ → 3′ order. The expected product length was listed as 524 base pairs.

The following table lists additional information regarding the primers designed, including length, melting and annealing temperatures (T_m_ and T_a_), GC content, and folding T_m_.

**Table 1:**
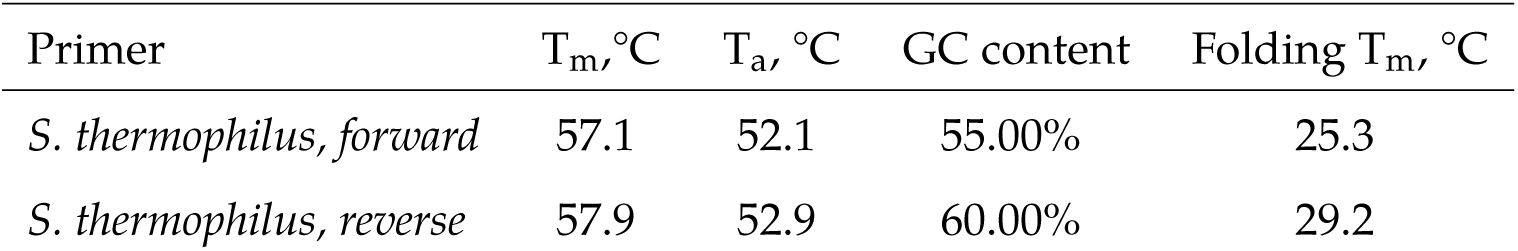
Primer Data. This table details pertinent primer design data for the primers used in this experiment, both forward and reverse.

| Primer | $T_m$, °C | $T_a$, °C | GC content | Folding $T_m$, °C |
| --- | --- | --- | --- | --- |
| *S. thermophilus*, forward | 57.1 | 52.1 | 55.00% | 25.3 |
| *S. thermophilus*, reverse | 57.9 | 52.9 | 60.00% | 29.2 |

Primers were resuspended from the lyophilized form in which they were delivered by the addition of molecular grade water to create a working stock; 275 μL of water was added for the forward primer, and 283 μL of water was added for the reverse primer.

### 2.5. PCR purification, gel electrophoresis, sequencing

All of these steps were performed by an external genomics service/CRO, Genewiz, LLC in South Plainfield, New Jersey. PCR products were purified by magnetic bead-based clean up, and were sequenced with standard Sanger sequencing methodologies.

### 2.6. Sequence processing

In order to produce FASTA files from.ab1 files (files containing electropherograms and base sequences), the program SnapGene Viewer was used. The FASTA files produced from the electropherograms were then processed through both the publicly available National Center for Biotechnology Information (NCBI) and Kyoto Encyclopedia of Genes and Genomes (KEGG) databases through their respective BLAST tools in order to identify and verify the identity of the sequenced DNA.

## 3. Results

### 3.1. Gel electrophoresis results

The results for gel electrophoresis are located in Figure 1. The figures are labeled with a short-hand labelling convention; this is in the form of [brand name], [first letter of bacterial genus tested]. Each brand name is also abbreviated; Chobani is CHOB, Greek Gods Greek Yogurt is GGGY,Yoplait is YPLT, and Fage remains FAGE. Unlabeled wells indicate results which do not belong to this experiment. For *S. thermophilus*, Chobani, Greek Gods Yogurt, and Fage give positive gel results, while Yoplait gives a negative result.

**Figure 1:**
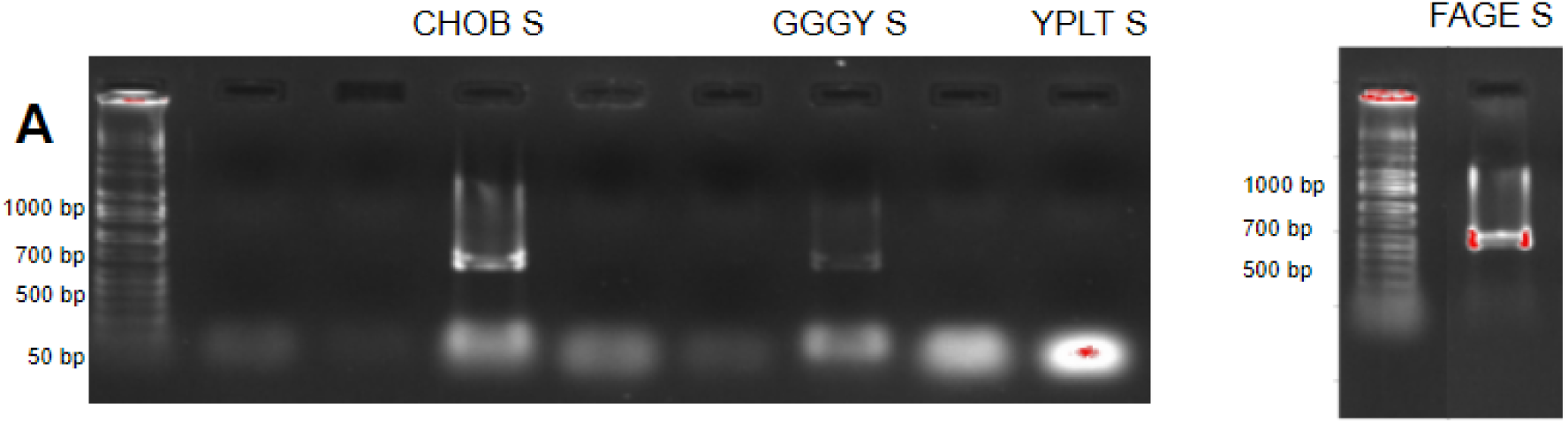
Annotated electrophoresis results. Row **A** contains all the electrophoresis results for *S. thermophilus*. CHOB S, GGGY S, and FAGE S were positive and are marked by red boxes. The e-gel DNA ladder from each gel is labeled with rough indicators of amplicon size in number of base pairs (bp). The results for FAGE S are shown seperately as they were processed on a separate gel.

The samples provided varying degrees of primer hybridization, especially in samples that provided visually weaker results. Stronger results are indicated by both stronger (whiter) bands in the gel result, and red marks indicate results with greater concentrations of DNA. This suggests greater concentrations of DNA product in the FAGE S sample, followed by the CHOB S and GGGY S samples respectively.

### 3.2. BLAST results (sequence identification)

After processing with publicly available BLAST tools from both NCBI and KEGG resources, all successful amplifications were identified as matching with other existing accessioned *S. thermophilus* samples. Table **??** lists a compiled record of accession numbers and match information for the primary matches of each sample sequenced and referenced against NCBI and KEGG databases. This data is accurate as of March 3rd, 2018.

The products of the Fage and Chobani samples all fully match to the same genome sample for *S. thermophilus* in the NCBI database, as indicated by a 100 percent identity percentage match to accession CP025400.1. The products of the Greek Gods Greek Yogurt sample indicate a 99 percent identity percentage match in the NCBI database to accession MG825731.1 (a partial sequence of *S. thermophilus* from the 16S-rRNA region of a bacterial sample).

## 4. Discussion of results

### 4.1. Result accuracy

#### Yoplait sample

The negative result for the Yoplait sample may be interpreted as no bacteria present in the sample; it may also be interpreted as being a false-negative due to a lack of bacterial DNA in the particular sample taken, and not in the yogurt itself. There is also a potential for false-negatives due to poor reaction performance; however, this might not be the case as the protocol followed is the same for all samples taken (3 of which were successful) and is based on a proven protocol for the detection of eDNA. There may very well have been no detectable bacterial DNA in the yogurt; evidently enough, the primers simply reproduced themselves (resulting in a strong band near the bottom of the gel image). While the yogurt claimed to include live and active cultures, no specifics were listed in the ingredients list on the yogurt tested itself.

#### Primers and their role in testing

The primers seem to have been non-specific enough to reproduce part of the 16S rRNA gene in one of the samples, which was not necessarily the original target sequence of the primers when they were created. At the very least, this could mean that the primers are not specific to a particular commercial strain of *S. thermophilus*, meaning that they can be used to amplify more strains and can be used on a wider range of commercially available commercial yogurt. However, they are are non-specific enough to reproduce at least two differing regions of the same bacterial genome.

### 4.2. Contaminants and culture amounts from the point of production

#### Yogurt contents and detection

It seems that all yogurts are simply not made equal. Commercially sold dietary yogurt products could variably contain substances that make it harder to determine the presence/absence of their claimed bacterial cultures.

The strongest bands and consistent results come from yogurts that only list two sets of ingredients; the Fage and Chobani-branded plain yogurt products tested sampled only list milk and yogurt cultures as their ingredients. Greek Gods Greek Yogurt, the one yogurt with a smaller match percentage to a more general gene of the bacteria, contained cream and pectin in addition to milk and yogurt cultures. These were most likely placed in the yogurt as thickeners.

The one negative result comes from Yoplait yogurt, which most prominently of various thickeners (such as pectin, gelatin, starches, corn syrups,and other oils), flavorings, and colorants. The yogurt tested was Yoplait Original Strawberry; while it claimed to contain live cultures, these were not explicitly listed on the ingredient list.

It is very likely that the presence of these other substances may cause issues in the detection of bacterial DNA, whether or not those bacteria are actually dead or living. Issues of contamination could also be the cause of the weaker bands in the Yoplait and Greek Gods samples; both of these samples have more ingredients that could dilute the presence of these bacterial cultures in yogurt.

## 5. Conclusions

### 5.1. Re-evaluating links between yogurt, probiotics, and health impacts

The uneven set of results raises concerns over the importance of bacterial concentration and the actual composition of yogurt biota on the effects of dietary yogurt. If yogurt with almost undetectable amounts of DNA present for probiotic strains of bacteria is credited for providing positive health impacts, are the bacteria themselves the beneficial agents making yogurt a preventative for disease? Or, are bacterial byproducts (the fermented yogurt itself) the actual source of these positive health impacts?

DNA testing through this cost-effective method could provide a goldstandard for bacterial identification for the biota present in dietary products. With DNA quantification methods, one could further determine the actual concentration and quality of DNA present in each sample mixture.

However, it might be more useful in certain cases to pursue further research through cell culturing of yogurt products to both provide a more complete picture of a commercial yogurt’s biome and make more generalized statements about the realities of the probiotic value of commercially sold yogurt. The contaminants which we speculated over may affect DNA testing for bacteria, but not culturing; we can attempt to determine the actual rate at which probiotics are present in dietary yogurt in order to determine whether it is the probiotics being live in the yogurt sold or simply the product of fermentative processing (yogurt itself, with or without live cultures) that provides these speculated benefits.

Specialized testing through DNA methods is most useful to come to important research leads about the causal link between bacteria and these positive health effects. More extensive testing on larger batches of yogurt could provide a better idea of overall product quality and consistency, leading to a baseline for consideration by nutritional and health researchers to further specify proven causal relationships between probiotics and human health benefits. The low cost and time required to prepare and process samples from the point of collection could also promote more consumer-oriented research into the quality of products sold; and, if this technology indeed is used to prove that there is a strong causal link between *S. thermophilus* and human health effects, could inform the creation of actionable points to reinforce recommendations on issues of public and preventative health, and could also lead to better decisions for consumers who try to take advantage of probiotics in products sold to them.

## Supporting information

## Data availability

All the data related to this set of experiments is included in this article as part of the text, as figures, or in the supporting information provided in the text. Additional supplementary files analyzed in this set of experiments have also been released.

## Supporting Information

### S1: FASTA file, FAGE S

This is a copyable version of the FASTA file for FAGE S which was interpreted from a chromatogram and used for BLAST.

>FAGE SAMPLE STREPTOCOCCUS THERMOPHILUS

AACGCATTAAGCACTCCGCCTGGGGAGTACGACCGCAAGGTTGAAACTCAAAGGAATTGACGGGGGCCCGCACAAGCGGTGGAGCATGTGGTTTAATTCGAAGCAACGCGAAGAACCTTACCAGGTCTTGACATCCCGATGCTATTTCTAAAGATAGAAAGTTACTTCGGTACATCGGTGACAGGTGGTGCATGGTTGTCGTCAGCTCGTGTCGTGAGATGTTGGGTTAAGTCCCGCAACGAGCGCAACCCCTATTGTTAGTTGCCATCATTCAGTTGGGCACTCTAGCGAGACTGCCGGTAATAAACCGGAGGAAGGTGGGGATGACGTCAAATCATCATGCCCCTTATGACCTGGGCTACACACGTGCTACAATGGTTGGTACAACGAGTTGCGAGTCGGTGACGGCGAGCTAATCTCTTAAAGCCAATCTCAGTTCGGATTGTAGGCTGCAACTCGCCTACATGAAGTCGGAATCGCTAGTAATCGCGGATCAGCACGCCGCGGTGAATACGTTCCCGGGCCTTGTACACACCGCCCGTCACACCACNAGAGTTTGTAACACCCGAAGTCGGTGAGGTAACCTTTTGGAGCCAGCCGCCTAAGGTGGGACANATGATTGGGGTGAAGTCGTAACAAGG

### S2: FASTA file, CHOB S

This is a copyable version of the FASTA file for CHOB S which was interpreted from a chromatogram and used for BLAST.

>CHOB SAMPLE STREPTOCOCCUS THERMOPHILUS

AAGGTTGAAACTCAAAGGAATTGACGGGGGCCCGCACAAGCGGTGGAGCATGTGGTTTAATTCGAAGCAACGCGAAGAACCTTACCAGGTCTTGACATCCCGATGCTATTTCTAGAGATAGAAAGTTACTTCGGTACATCGGTGACAGGTGGTGCATGGTTGTCGTCAGCTCGTGTCGTGAGATGTTGGGTTAAGTCCCGCAACGAGCGCAACCCCTATTGTTAGTTGCCATCATTCAGTTGGGCACTCTAGCGAGACTGCCGGTAATAAACCGGAGGAAGGTGGGGATGACGTCAAATCATCATGCCCCTTATGACCTGGGCTACACACGTGCTACAATGGTTGGTACAACGAGTTGCGAGTCGGTGACGGCGAGCTAATCTCTTAAAGCCAATCTCAGTTCGGATTGTAGGCTGCAACTCGCCTACATGAAGTCGGAATCGCTAGTAATCGCGGATCAGCACGCCGCGGTGAATACGTTCCCGGGCCTTGTACACACCGCCCGTCACACCACGAGAGTTTGTAACACCCGAAGTCGGTGAGGTAACCTTTTGGAGCCAGCCGCCTAAGGTGGGACAGATGATTGGGGTGAAGTCGTAACAAG

### S3: FASTA file, GGGY S

This is a copyable version of the FASTA file for GGGY S which was interpreted from a chromatogram and used for BLAST.

>GGGY SAMPLE STREPTOCOCCUS THERMOPHILUS

ACGACCGCAAGGTTGAAACTCAAAGGAATTGACGGGGGCCCGCACAAGCGGTGGAGCATGTGGTTTAATTCGAAGCAACGCGAAGAACCTTACCAGGTCTTGACATCCCGATGCTATTTCTAAAAATAGAAAGTTACTTCGGTACNTCGGTGACAGGTGGTGCATGGTTGTCGTCAGCTCGTGTCGTGAGATGTTGGGTTAAGTCCCGCAACGAGCGCAACCCCTATTGTTAGTTGCCATCATTCAGTTGGGCACTCTAACGAGACTGCCGGTAATAAACCGGAGGAAGGTGGGGATGACGTCAAATCATCATGCCCCTTATGACCTGGGCTACACACGTGCTACAATGGTTGGTACAACGAGTTGCGAGTCGGTGACGGCGAGCTAATCTCTTAAAGCCNATCTCAGTTCGGATTGTAGGCTGCAACTCGCCTACATGAAGTCGGAATCGCTAGTAATCGCGGATCAGCACGCCGCGGTGAATACGTTCCCGGGCCTTGTACACACCGCCCGTCACACCNNNAGAGTTTGTANNNNNCGAAGTCGGTGAGGTAACCTTTTGGAGCCAGCCGCCTAAGGTGGGACAGATGATTGGGGTGAAGTCGTAACA

### S4: .zip archive, chroma_STherm.zip

This archive contains the chromatogram files for each sample tested.

The following is a list of the zip-file’s contents.

*FAGE_S-Stherm.ab1.* A file containing a full chromatogram for the Fage sample, which can be viewed using many different programs dedicated to viewing chromatograms and can also be converted to FASTA files as was done in this paper.
*CHOB_S-Stherm.ab1.* A file containing a full chromatogram for the Chobani sample.
*GGGY_S-Stherm-ab1.* A file containing a full chromatogram for the Greek Gods Greek Yogurt sample.

